# Natural monoclonal autoantibodies against HERV-K102 Envelope-TM from SLE patients selectively eliminate autoreactive immune cells and cancer cells

**DOI:** 10.1101/2024.12.14.628522

**Authors:** Qinyuan Gong, Mengyuan Li, Shuwen Zheng, Zhaoxing Wu, Ping Wang, Xuzhao Zhang, Yun Liang, Wenbin Qian, Rongzhen Xu

**Affiliations:** Department of Hematology and Cancer Institute (Key Laboratory of Cancer Prevention and Intervention, China National Ministry of Education), the Second Affiliated Hospital, Zhejiang University School of Medicine, Hangzhou 310009, China; Institute of Hematology, Zhejiang University, Hangzhou, 310009, China

**Keywords:** endogenous retroviruses, HERV-K102, Systemic lupus erythematosus, cancer, monoclonal autoantibodies

## Abstract

In sharp contrast to immune dysfunction in cancer, autoimmune diseases such as systemic lupus erythematosus (SLE) exhibit excessive immune reactions characterized by high titers of autoantibodies to HERV-K102-envelope (HERV-K102-Env) neoantigens, which are frequently found in cancer patients but seldom elicit a strong immune response. Here we aim to address whether anti-HERV-K102-Env autoantibodies in SLE can effectively eliminate autoreactive immune cells and cancer cells expressing HERV-K102-Env neoantigens. We established world-first fully human autoantibody phage display library with 3.67x10^8^ cfu using peripheral blood mononuclear cells (PBMCs) from SLE patients. Through high-throughput screening, we identified nineteen monoclonal autoantibodies (mAbs) targeting the conserved HERV-K102 Env-TM subunit. The EC50 values of these autoantibodies binding to the HERV-K102 Env-TM subunit ranged from 0.002436 μg/ml to 1.798 μg/ml. Remarkably, eleven of these mAbs not only recognized the HERV-K102 Env-TM glycoprotein on the cell surface but also effectively eliminated autoreactive B, T, and natural killer (NK) cells in SLE, as well as cancer cells. Our findings provide a conceptually new immunotherapy target HERV-K102 Env-TM subunit and open the era of using natural human autoantibodies to treat autoimmune disease and cancers.

Several of these monoclonal autoantibodies show promise as potential diagnostic and therapeutic agents, paving the way for innovative approaches to treating SLE and various malignancies.

## Introduction

Immunotherapy has revolutionized cancer treatment. For instance, immune checkpoint inhibitors (ICIs) –based immunotherapies, such as anti-PD1/anti-PDL1, have led to durable remissions in a variety of different tumor types^1–3^. CAR-T cell immunotherapies have proven effective in patients with leukemia, B cell lymphoma, and multiple myeloma (MM)^4–6^, as well as systemic lupus erythematosus (SLE)^7–9^. However, most cancer patients remain refractory to current immunotherapeutic approaches, necessitating the development of novel strategies to overcome resistance^10–15^.

Immune dysfunction and excessive immune reactions are well-established hallmarks of cancer^10–15^ and SLE^16–19^, respectively. However, there is currently no information available regarding the potential relationship between cancer and autoimmune diseases. Previous research has demonstrated that neoantigens derived from human endogenous retroviruses (HERVs), such as HERV-K102 envelope, are frequently enriched in cancer tissues across multiple cancer types, including acute myelogenous leukemia (AML), pancreatic ductal adenocarcinoma (PDAC), lung cancer, and glioma^20–25^. Conversely, these neoantigens rarely elicit a strong immune response in cancer patients. In contrast, autoantibodies against HERV-K102 envelopes are markedly increased in SLE patients^16,17^. This paradoxical phenomenon prompted us to investigate whether autoantibodies against HERV-K102 envelope in SLE patients could eliminate autoreactive immune cells in autoimmune diseases and cancer cells.

In this study, we first attempted to construct a fully human HERV-K102 Env-TM autoantibody phage display library from PBMCs of SLE patients and then identified monoclonal autoantibodies through high-throughput screening with HERV-K102-TM or SU antigens, and finally evaluate the potential diagnostic and therapeutic values of these monoclonal autoantibodies in SLE and cancers.

## Materials and Methods

### Collection of Patient Serum Samples

This study was performed after approval by the Ethics and Scientific Committee of The Second Affiliated Hospital, Zhejiang University School of Medicine (No. 2024-0325). To characterize the autoantibody expression profile of HERV-K102 Env-TM in patients with autoimmune disease and cancer, serum samples were collected from thirteen SLE patients and fifteen PDAC patients using a standardized protocol and stored at -80°C until analysis. The serum samples from all participants were evaluated for titers of autoantibodies against the HERV-K102 Env-TM antigen.

### ELISA for detection of serum autoantibodies against HERV-K102 Env-TM antigen

Serum autoantibody titers against HERV-Env-TM were determined using an enzyme-linked immunosorbent assay (ELISA) with purified recombinant HERV-K102 Env-TM glycoprotein produced in our laboratory. Briefly, a 96-well ELISA plate was coated with HERV-K102 Env-TM proteins (4 μg/ml) and blocked with 5% phosphate-buffered saline containing milk (PBSM). Sera from SLE and PDAC patients were added to the coated wells in serial dilutions (1:100-1:256,000) with phosphate-buffered saline (PBS), followed by detection using horseradish peroxidase (HRP)-conjugated anti-human IgG antibodies.

### Phage display library construction of PBMCs from SLE patients

PBMCs were isolated from the blood samples of 32 SLE patients with high titers of autoantibodies to HERV-K102 Env-TM antigen. Total RNA was extracted and used to synthesize cDNA through reverse transcription. PCR was employed to amplify the genes encoding the heavy and light chains of the antibody. The full-length coding sequence was obtained by linking the open reading frame (ORF) of light and heavy chain genes in vitro. The vector and the full-length coding sequence were separately digested, and the antibody nucleotide sequence was cloned into the *M13* filamentous phage expression vector. The construct was then transferred into *E. coli TG1* competent cells via electrotransformation, followed by the addition of helper phage *M13K07* to facilitate phage proliferation and display antibody expression on the surface.

The phage antibody library was obtained by centrifuging the bacteria and precipitating phages from the supernatant. The dilution spot plate method showed that the capacity of this antibody library was 3.67×10^8^ cfu. Sequencing analysis revealed a correct insertion rate exceeding 80%, and the high abundance was reflected by the CDR3 region.

### Screening and characterization of autoantibodies against K102-Env-TM or SU antigens

Phage display technology was employed to display the constructed antibody sequences of the full human antibody library on the phage surface. Specific proteins, cells or peptides were used as antigens, and specific monoclonal antibodies against the antigens were enriched through several rounds of selection. Subsequently, positive clones that specifically bind with the antigens were screened by ELISA, and sequencing was performed to obtain the antibody sequences of these positive clones. From the enriched pool, a total of 512 monoclones were selected for screening, and 101 positive clones bound to the HERV-K102-TM antigen and 48 positive clones bound to the HERV-K102-SU antigen were obtained. Sequencing analysis was conducted on 135 of these clones, resulting in the identification of 68 unique molecules. Twenty of these molecules were selected for full-length construction, while the remaining molecules were excluded from full-length construction due to the presence of post-translational modification sites or limited diversity.

### Binding assay of autoantibodies to HERV-K102-TM or SU antigens by ELISA

By constructing a phage display library, we screened a series of monoclonal autoantibodies exhibiting high specificity and strong binding affinity to HERV-K102-TM or HERV-K102-SU antigens. Competitive effects of 20 monoclonal antibodies binding to the HERV-K102-TM antigen with the positive control biotin-conjugated monoclonal antibody (1G6-Biotin) were determined by ELISA. ELISA plates were coated with screened antibodies at a concentration of 2 μg/mL in 1×PBS, 30 µL per well, and incubated overnight at 4°C. The following day, plates were blocked with 5% PBSM for 2 hours at room temperature (RT). Diluted K102-TM-hFc antigen in 1% PBSM was added to the ELISA plates, 30 µL per well, and incubated for 60 minutes at RT. Subsequently, 1G6-Biotin antibody against HERV-K102 Env-TM antigen in 1% PBSM was added, 30 µL per well, for 60 minutes at RT. Neutravidin-HRP (Thermo Fisher, 31001), diluted 1:2000 in 1×PBS, was added at 30 µL per well and incubated for 50 minutes at RT. Finally, Tetramethylbenzidine (TMB) substrate was added, and the reaction was terminated with 2M stop solution. The optical density at 450 nm (OD450) was measured.

The affinities of 20 monoclonal antibodies to HERV-K102 SU antigen were determined by ELISA. The ELISA plates were coated with K102-SU-FC antigen at a concentration of 2 μg/mL in 1× PBS, using 30 µL per well, and incubated overnight at 4°C. The following day, the plates were blocked with 5% PBSM for 2 hours at RT. Diluted antibodies in 1% PBSM were added to the ELISA plates at 30 µL per well and incubated for 60 minutes at RT. Subsequently, Anti-human-κ+λ-HRP (Millipore, AP502P, AP506P) was added at a 1:4000 dilution in 1% PBSM, using 30 µL per well, and incubated for 50 minutes at RT. Finally, TMB substrate was added, and the reaction was stopped with 2M stop solution. The OD450 was measured.

### Cell Lines and Culture

This study employed a panel of 10 diverse human tumor cell lines, all cultured in media supplemented with 10% fetal bovine serum (FBS) and 1% penicillin-streptomycin. The AML cell lines MOLM-13, Kasumi-1, HL-60, NB4, THP-1, OCI-AML3 and MV4-11 were cultured in RPMI-1640 medium, while KG-1a was cultured in IMDM. The human solid tumor cell lines MIA-PaCa2 and PANC-1, as well as the murine cell line CHO, were cultured in DMEM. All cell lines were cultured in a humidified atmosphere containing 5% CO_2_ at 37°C and routinely screened for mycoplasma contamination. Experiments were conducted exclusively with cells confirmed to be free of mycoplasma.

### Western blotting analysis and Antibodies

To determine protein expression levels by Western blotting, cells were collected and washed twice with pre-cooled PBS (Cinery). M-PER Mammalian Protein Extraction Reagent (Thermo, 78501) containing 1% protease and phosphatase inhibitor cocktail (Thermo, 78428) was used for cell lysis on ice for 30 min. Cell lysates were then centrifuged at 13,000×g for 15 min, and supernatants were collected and heated to 100°C for 10 min. Protein concentrations were quantified using the Pierce BCA Protein Assay Kit (Thermo, 23225). Equal quantities of proteins were subjected to Sodium Dodecyl Sulfate-Polyacrylamide Gel Electrophoresis (SDS-PAGE) and transferred to Polyvinylidene Fluoride (PVDF) membranes (BIO-RAD, #1620177). Then the membranes were blocked with 5% nonfat milk and incubated with primary antibodies (diluted serum of patients or HERV-K102 Env-TM autoantibodies). After overnight incubation at 4°C, membranes were washed three times and probed with horseradish peroxidase-conjugated secondary antibodies (goat anti-rabbit (Huabio, HA1001), goat anti-mouse (Huabio, HA1006), and Goat Anti-Human IgG Fc (Abcam, AB97225)) for 1 hour at RT. Signals were detected using the Tanon 5200 Chemiluminescent Imaging System.

### Detection of HERV-K-TM autoantibody-reactive antigens on immune cells in SLE and primary AML cancer cells

Primary peripheral blood samples were obtained from clinical residual blood samples of 17 healthy individuals, 10 SLE patients, and 5 patients with primary AML. PBMCs, which contain immune cells or primary AML cells, were isolated through density gradient centrifugation with Lymphocyte Separation Medium (TBD). After centrifugation, the white membrane layer in the middle of the sample was collected and washed with PBS.

### Flow cytometry (FCM)

FCM was performed using a NovoCyte flow cytometer (ACEA Biosciences). AML cell lines and PBMCs from healthy individuals and AML patients were stained with HERV-K102 Env-TM autoantibody and the secondary antibody PE F(ab’)2 Goat anti-human IgG Fcγ (BioLegend, Cat 398004), as well as PE Mouse IgG1, κ Isotype Ctrl (FC) (Cat 400113). PBMCs from SLE patients were stained with mouse anti-human monoclonal antibodies purchased from BioLegend, including PB CD45 (Cat 304022), FITC CD3 (Cat 317306), APC CD3 (Cat 300412), CD19-APC (Cat 302211), CD56-APC/Cyanine 7 (Cat 318331), and APC Granzyme B (Cat 372203), in addition to the HERV-K102 Env-TM antibody. PD-1-PE/Cy7 (Cat 561272) was obtained from BD Biosciences. The IFN-α antibody (sc-373757) was acquired from Santa Cruz, and the secondary antibody FITC-AffiniPure Goat Anti-Mouse IgG(H+L) (33207ES60) was purchased from Yeasen. After cell membrane permeabilization, intracellular IFN-α expression in T cells and granzyme B (GZMB) expression in NK cells were investigated.

### Immunofluorescent staining for the distribution of HERV-K102-TM autoantibody-reactive antigens in cancer cells

AML and PDAC cells were seeded on coverslips and fixed in 4% paraformaldehyde, then blocked with 10% rabbit serum. The coverslips were incubated with the monoclonal autoantibody 186 as the primary antibody in blocking buffer at 4°C overnight, followed by incubation with the secondary antibody PE F(ab’)2 Goat anti-human IgG Fcγ (BioLegend, Cat 398004) for 1h at room temperature. After washing off excess secondary antibodies, cell nuclei were stained using 4’,6-diamidino-2-phenylindole (DAPI). The slides were imaged using a Zeiss Confocal Laser Scanning Microscope 710 (LSM710, Germany). The ZEN software was used for image acquisition and analysis.

### Antibody-dependent Cell-mediated Cytotoxicity (ADCC) assay

PBMCs were isolated from healthy individuals as effector cells and cultured with MIA-PaCa2 cells or OCI-AML3 cells with luciferin tag (target cells) at an effector-to-target (E:T) ratio of 3:1. MIA-PaCa2 cells were pre-seeded into 96-well plates at a density of 4,000 cells per well, while OCI-AML3 cells were plated at a density of 10,000 cells per well. HERV-K102 Env-TM autoantibodies at various concentrations (1 μg/mL, 5 μg/mL) were added to mediate the killing effect, compared to a control group without autoantibody.

After 24 hours of incubation, fluorescent intensity was detected at a wavelength of 562 nm, and the remaining surviving MIA-PaCa2 or OCI-AML3 cells were quantified. PBMCs isolated from SLE patients were equally divided into five parts. The experimental group was incubated with monoclonal autoantibody 910 at different concentrations (0.1 μg/mL, 0.5 μg/mL, 1 μg/mL) compared to a control group without monoclonal autoantibody 910 in IMDM for 24 hours. The target cells were HERV-K-TM^+^ immune cells, and the effector cells were those acting as killers in PBMCs from the same blood sample. FC was used to detect positive rates of K-TM^+^ cells and total immune cells before and after co-incubation.

### Statistical analysis

All statistical analyses were performed using GraphPad Prism 9.0 and IBM SPSS. Parametric comparisons of normally distributed values meeting the variance test were conducted using an unpaired Student’s t-test, while data failing to meet the variance test were compared using the Wilcoxon rank sum test. A p-value less than 0.05 was considered statistically significant, and significance levels were indicated as follows: * p < 0.05, ** p < 0.01, *** p < 0.001, and ns p > 0.05.

## Results

### SLE patients possess high titers of autoantibodies against HERV-K102-Env-TM antigens

The envelope protein of HERV-K (HERV-K Env) consists of a surface subunit (SU) and a transmembrane subunit (TM)^26^. Previous studies have demonstrated the frequent presence of autoantibodies targeting SU antigens in the sera of SLE patients^16,17^. However, the presence of autoantibodies specifically targeting HERV-K102 Env-TM antigens in the blood of SLE patients remains unexplored. To elucidate the expression levels of anti-HERV-K102 Env-TM autoantibodies in patients with autoimmune diseases and cancer, we employed ELISA to examine the titers of autoantibodies targeting the K102-Env-TM antigen in sera from SLE and PDAC patients. Our findings revealed significantly higher titers of anti-K-TM antibodies in the sera of SLE patients compared to PDAC patients (Fig. 1a). The titers of anti-K-TM autoantibodies in SLE sera exhibited substantial variability, ranging from 1:1,000 to 1:256,000, with some SLE cases displaying titers as high as 1:256,000. In contrast, the titers of anti-K-TM autoantibodies were low or undetectable in the sera of PDAC patients. These results were further corroborated by Western blotting, with three representative samples from SLE and PDAC patients shown in Fig.1b. This intriguing paradox motivated our investigation into the potential anti-autoimmune disease and anti-tumor activity of these autoantibodies targeting the HERV-K102 Env-TM antigen in SLE patients.

**Fig. 1.**
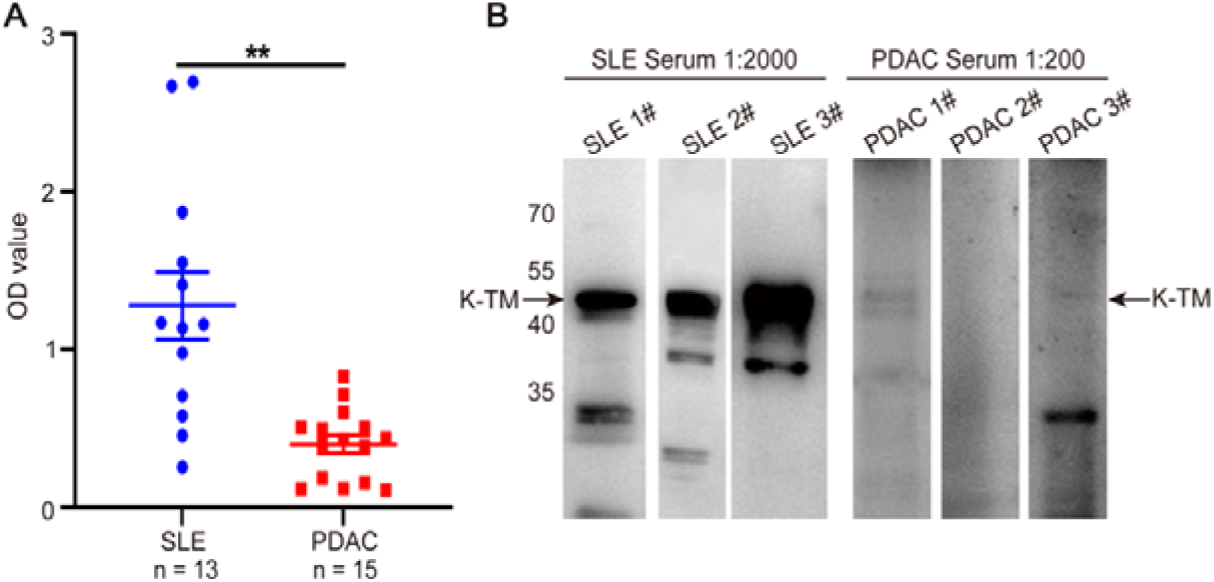
K102-Env-TM autoantibodies are markedly increased in SLE sera but low or absent in PDAC sera. (**A**) The titers of autoantibodies targeting the K102-Env-TM antigen in sera of SLE and PDAC patients were determined by ELISA. (**B**) K-TM recombinant proteins were incubated with diluted serum from three representative SLE patients (1:2000) and three representative PDAC patients (1:200) by Western blotting.

### Phage display library construction and characterization of human monoclonal autoantibodies to HERV-K102 Env-TM and SU antigens

Based on our findings that SLE patients exhibit a high titer of autoantibodies to the HERV-K-TM antigen in their circulating blood, we hypothesized the presence of highly potent protective monoclonal autoantibodies against the HERV-K102 Env-TM antigen generated by B cells in SLE patients. Thus, we focused on identifying these putative autoantibodies. To achieve this, we first attempted to generate a phage display library from PBMCs of SLE patients with a high titer of autoantibodies to HERV-K102 Env-TM antigens.

Subsequently, we conducted a high-throughput screen with the HERV-K102 Env-TM glycoprotein and the K-SU glycoprotein.

A fully human autoantibody phage display library with 3.67×10^8^ cfu was successfully constructed. High-throughput screening with HERV-K102-TM or SU antigens yielded 101 K-TM antigen-binding IgG1^+^ clonotypes and 48 K-SU antigen-binding IgG1^+^ clonotypes. Sequencing analysis was performed on 135 selected clonotypes and we obtained 68 unique molecules, 20 of which were selected for full-length construction. The binding affinity of the candidate antibodies to K102-TM-hFc antigen, as determined by ELISA, is presented in Fig. 2A-C and Table 1-3. The EC50 values of these autoantibodies for binding the TM subunit ranged from 0.002436 μg/ml to 1.798 μg/ml. The binding affinity of the candidate antibodies to K102-SU-Fc antigen is depicted in Supplementary Fig. 1A, B and Supplementary Table 1, 2. Eleven mAb clones only reacted with K102-Env-TM glycoprotein, eight mAb clones reacted with both K-SU and K-TM, and one mAb clone only reacted with K-SU, as determined by ELISA binding assay (Supplementary Table3).

**Fig. 2.**
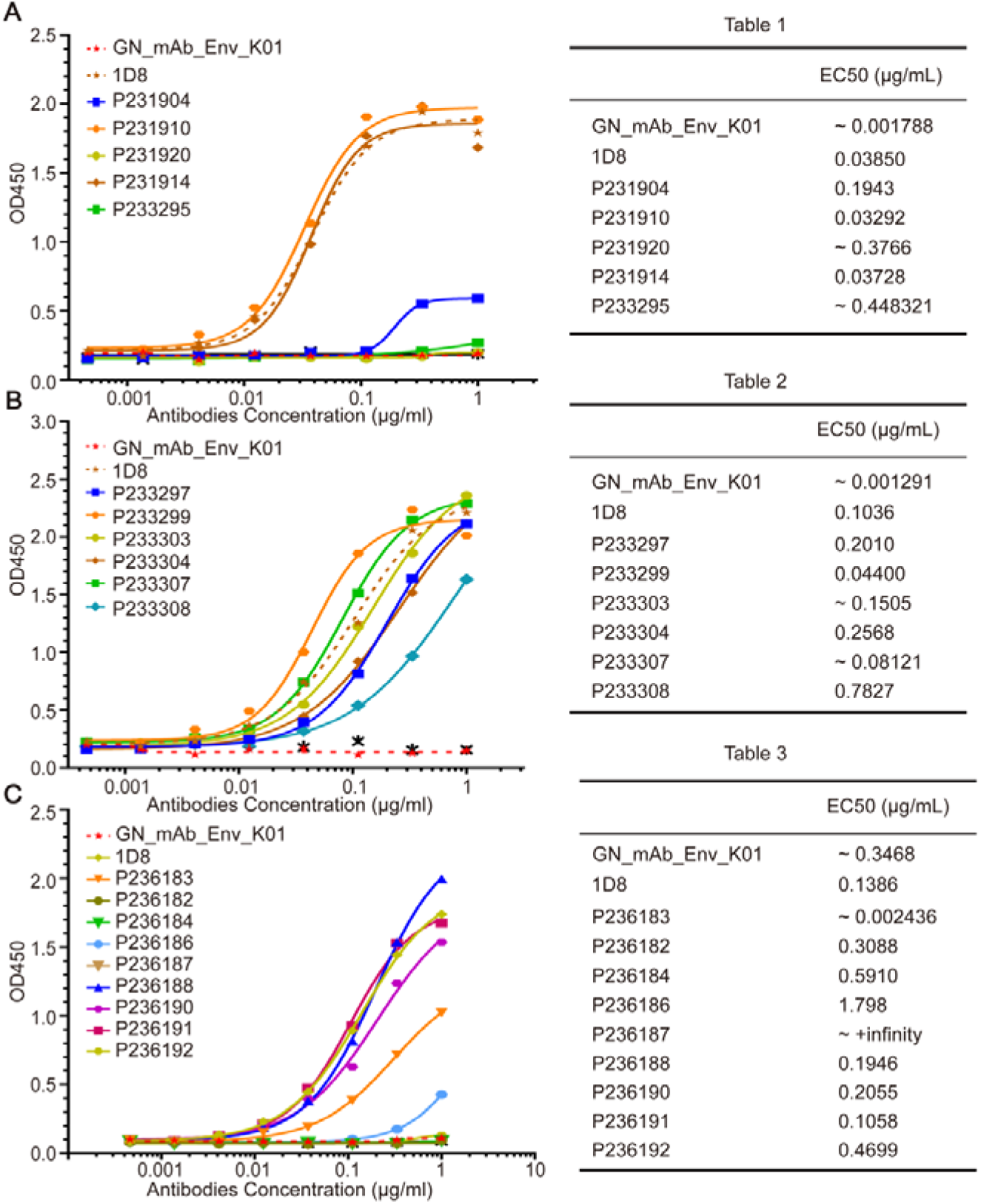
Competitive effects of 20 candidate antibodies binding to K102-TM-hFc antigen with positive control biotin-conjugated monoclonal antibody (1G6-Biotin) as determined by ELISA. (**A**) 904, 910, 920, 914, 295 antibodies. (**B**) 297, 299, 303, 304, 307, 308 antibodies. (**C**) 183, 182, 184, 186, 187, 188, 190, 191 and 192 antibodies. The positive control 1D8 antibody exhibits a high degree of binding to the K-TM antigen, while the positive control GN_mAb_Env_K01 antibody demonstrates a significant binding affinity for the K-SU antigen.

**Table 1.** Median effect concentration (EC50) values of 904, 910, 920, 914, 295 antibodies.

**Table 2.** EC50 values of 297, 299, 303, 304, 307, 308 antibodies.

**Table 3.** EC50 values of 183, 182, 184, 186, 187, 188, 190, 191 and 192 antibodies.

Taken together, we established the world-first phage display library of human autoantibodies against K102-Env-TM antigen s and identified a set of monoclonal autoantibodies with high specificity and potent binding ability with K-TM glycoprotein antigen.

### Active SLE patients exhibit elevated levels of HERV-K102 Env-TM^+^ immune cells

To determine whether HERV-K102 Env-TM autoantibody-reactive antigens are present in immune cells of SLE, we detected the proportion of HERV-K102 Env-TM^+^ T, B and NK cells in the blood of SLE patients with active disease.

Lymphocyte compartmentalization was shown with FCM plots and typical gating strategies. After gating on live cells and single cells, CD45^+^ leukocytes were selected, followed by CD3^+^ T cells, CD3^-^CD19^+^ B cells, and CD3^-^CD56^+^ NK cells, from which HERV-K102 Env-TM^+^ cells were identified. FCM results from 1# SLE patient revealed that the positive rate of K-TM^+^ T cells within the CD3^+^ T cell population was 31.45%, the positive rate of K-TM^+^ B cells within the CD19^+^ B cell population was 52.16%, and the positive rate of K-TM^+^ NK cells within the CD56^+^ NK cell population was 48.89% (Fig. 3A). In contrast, we also assessed the proportion of K-TM^+^ immune cells in the blood of healthy controls. FCM results from one control case, shown in Fig. 3B, and the positive rates of K-TM^+^ T, B and NK cells were 2.12%, 9.88%, and 10.76%, respectively. FCM plots of 2#, 3# SLE patients (A, B) and 2#, 3# healthy controls (C, D) are presented in Supplementary Fig. 2. K-TM^+^ T cells significantly increased in SLE patients (n = 8) compared to healthy controls (n = 8) (p < 0.001) (Fig. 3C). Similarly, K-TM^+^ B cells and K-TM^+^ NK cells were significantly elevated in SLE patients compared to healthy controls (both p < 0.001) (Fig. 3D, E).

**Fig. 3.**
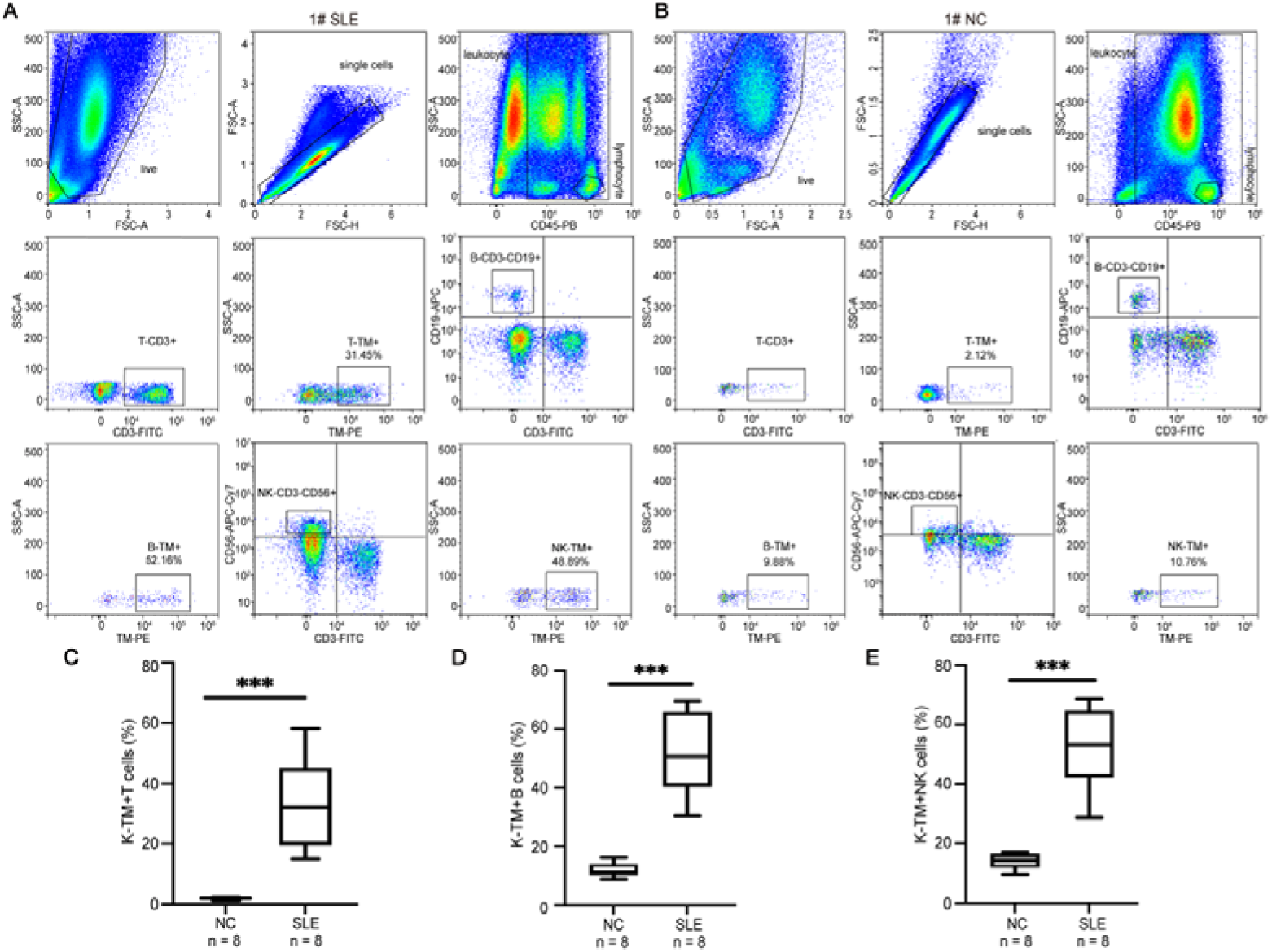
The proportions of HERV-K102 Env-TM^+^ T, B and NK cells in the blood samples of SLE patients compared to healthy controls. (**A**) Flow cytometry plots and gating strategies used to identify HERV-K102 Env-TM^+^ immune cells in 1# SLE patient. (**B**) FCM plots and gating strategies used to identify K-TM^+^ immune cells in 1# healthy control. (**C, D, E**) The proportions of K-TM^+^ T, B and NK cells in SLE patients (n = 8) and healthy controls (n = 8). *** p < 0.001.

### HERV-K102 Env-TM+ T cells and NK cells are autoreactive immune cells

The results demonstrated that active SLE patients exhibited elevated levels of HERV-K-TM^+^ immune cells, including B cells, T cells and NK cells, in comparison to healthy controls. IFN-α is recognized as a critical effector in both innate and adaptive immunity, contributing to the onset and progression of SLE^27–30^. While plasmacytoid dendritic cells (pDCs) are considered as the primary cell type producing high levels of IFN-α^31,32^, it remains unclear whether HERV-K102 Env-TM^+^ T cells also secrete IFN-α. Therefore, we used FCM to examine the proportion of IFN-α^+^ cells within the K102-Env-TM^+^ T cell population. We discovered that nearly all K102-Env-TM^+^ T cells were IFN-α^+^ cells in blood samples obtained from active SLE patients (Fig. 4A). The proportion of K-TM^+^ IFN-α^+^ T cells within the CD3^+^ T cell population was significantly increased in patients with active disease compared to healthy controls (p < 0.001) (Fig. 4B). Furthermore, HERV-K102 Env-TM^+^ NK cells from active SLE patients displayed elevated levels of GZMB, and the percentage of K-TM^+^ GZMB^+^ cells in NK cells of active SLE patients was higher than that of healthy controls (p < 0.001) (Fig. 4C). In summary, the HERV-K102 Env-TM^+^ T cells and NK cells might be pathogenic autoreactive immune cells in SLE.

**Fig. 4.**
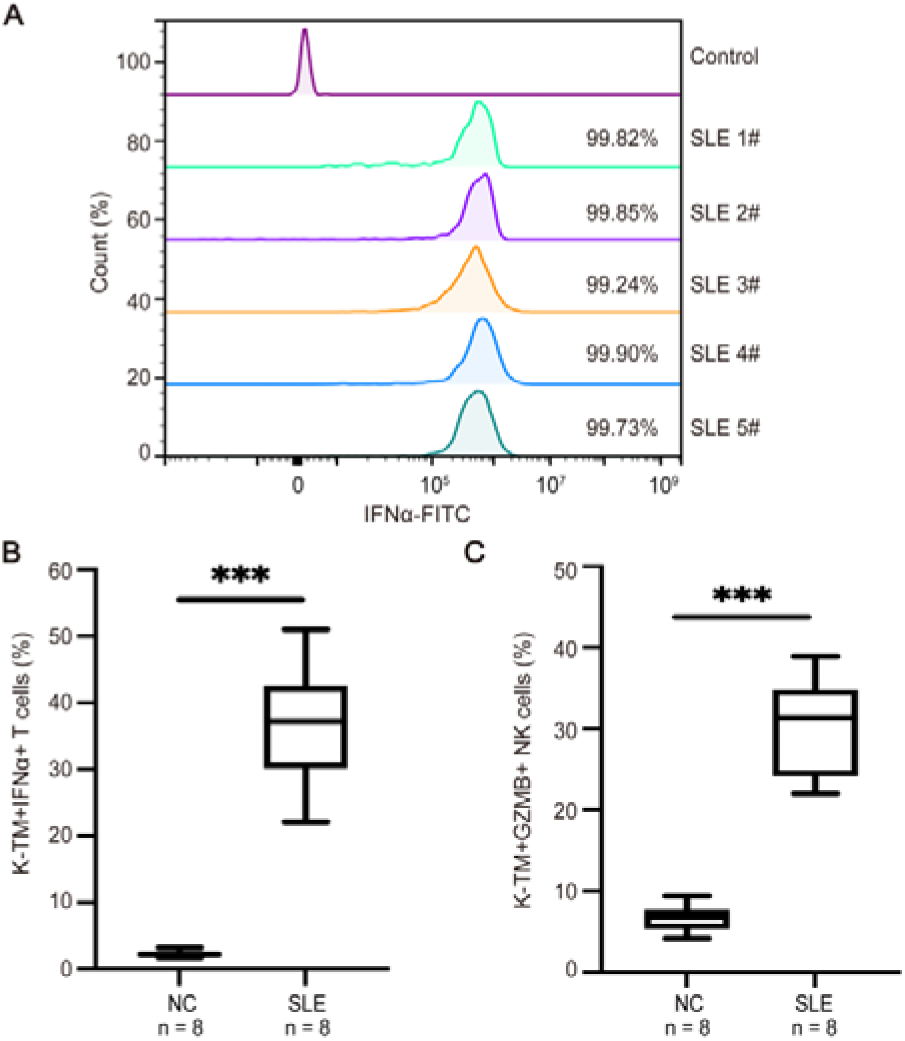
HERV-K102 Env-TM^+^ T cells and NK cells are autoreactive immune cells in SLE. **(A)** The proportion of IFN-α^+^ cells among K102-Env-TM^+^ T cells in active SLE patients (n = 5). **(B)** The proportion of K102-Env-TM^+^IFN-α^+^ T cells between patients with active SLE (n = 8) and healthy controls (n = 8). **(C)** The proportion of K102-Env-TM^+^GZMB^+^ NK cells between patients with active SLE (n = 8) and healthy controls (n = 8). *** p < 0.001.

### Monoclonal autoantibodies induce ADCC activity against HERV-K102

#### Env-TM^+^ immune cells in SLE

Our previous study demonstrated that most HERV-K102-Env-TM^+^ T cells are double positive for IFN-α and PD-1, suggesting their potential role in triggering T cell exhaustion. In addition, HERV-K102-Env-TM^+^ NK cells in SLE patients exhibited elevated levels of GZMB. We observed a high frequency of HERV-K102 Env-TM^+^ T, B, and NK cells in SLE patients, with these aberrant immune cells playing a crucial role in autoantibody and cytokine production, potentially serving as a critical etiology in SLE. To investigate whether HERV-K102 Env-TM autoantibodies could eliminate HERV-K-TM^+^ immune cells in SLE, we isolated PBMCs from two SLE patients and incubated them with monoclonal autoantibody 910 at various concentrations (0.1 μg/mL, 0.5 μg/mL and 1 μg/mL). FCM was used to detect the positive rates of subpopulations expressing HERV-K102 Env-TM antigens in total immune cells after incubation. Monoclonal autoantibody 910 treatment markedly reduced the proportions of HERV-K102 Env-TM^+^ T, B and NK cells. In 1# patients, the mean positive rates of HERV-K-TM^+^ T, B, and NK cells in total immune cells decreased to 9.89%, 11.22%, and 10.49%, respectively (Fig. 5A-C). Similarly, in 2# patient, the mean positive percentages decreased to 7.27%, 15.32% and 26.16%, respectively (Supplementary Fig. 3A-C). These results indicate that HERV-K102 Env-TM monoclonal autoantibody selectively eliminates HERV-K102 Env-TM^+^ immune cells while sparing HERV-K102 Env-TM^-^ cells, effectively normalizing immune cell subsets in the blood.

**Fig. 5.**
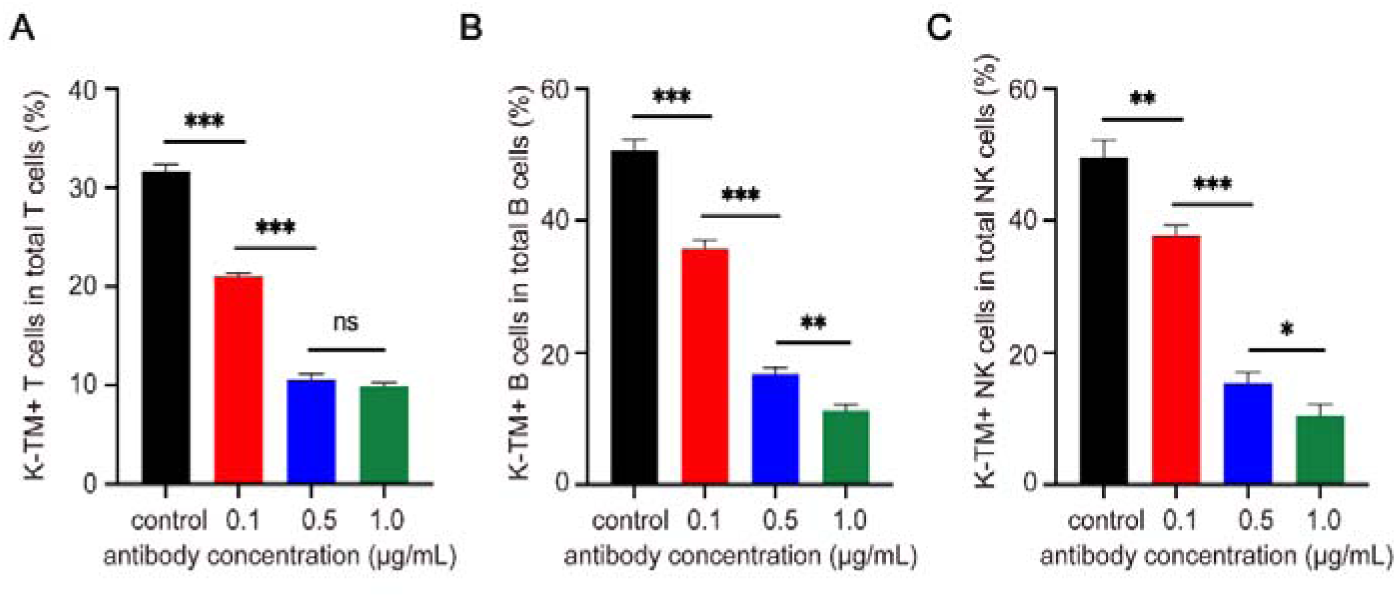
The ADCC activity of HERV-K102 Env-TM monoclonal autoantibody against HERV-K102 Env-TM^+^ T, B and NK cells in 1# patient with active SLE. (**A-C**) Various concentrations of the autoantibody (0.1 μg/mL, 0.5 μg/mL and 1 μg/mL) eliminated K-TM^+^ T, B, and NK cells from SLE patients. Data are presented as mean values ± s.d. of 3 biological replicates. * p < 0.05, ** p < 0.01, *** p < 0.001, and ns p > 0.05.

### Autoantibody-reactive HERV-K102 Env-TM antigens are abundantly present in cancer cells

To assess the presence of HERV-K102 Env-TM autoantibody-reactive antigens in cancer cells, the expression levels of these antigens were examined in PDAC cell lines and primary tumor specimens using Western blotting. K-TM autoantibody-reactive antigens were detected in the pancreatic cancer cell lines MIA-PaCa2 and Panc-1 but were absent in the CHO cell line (Fig. 6A). Subsequently, the expression of K-TM autoantibody-reactive antigens was investigated in 7 pairs of primary PDAC tumor and adjacent noncancerous tissues by WB using HERV-K102 Env-TM monoclonal autoantibody mAb 186. The results demonstrated that K-TM autoantibody-reactive antigens were highly expressed in all tested tumor tissues but were low or absent in the nonmalignant tissues (Fig. 6B).

**Fig. 6.**
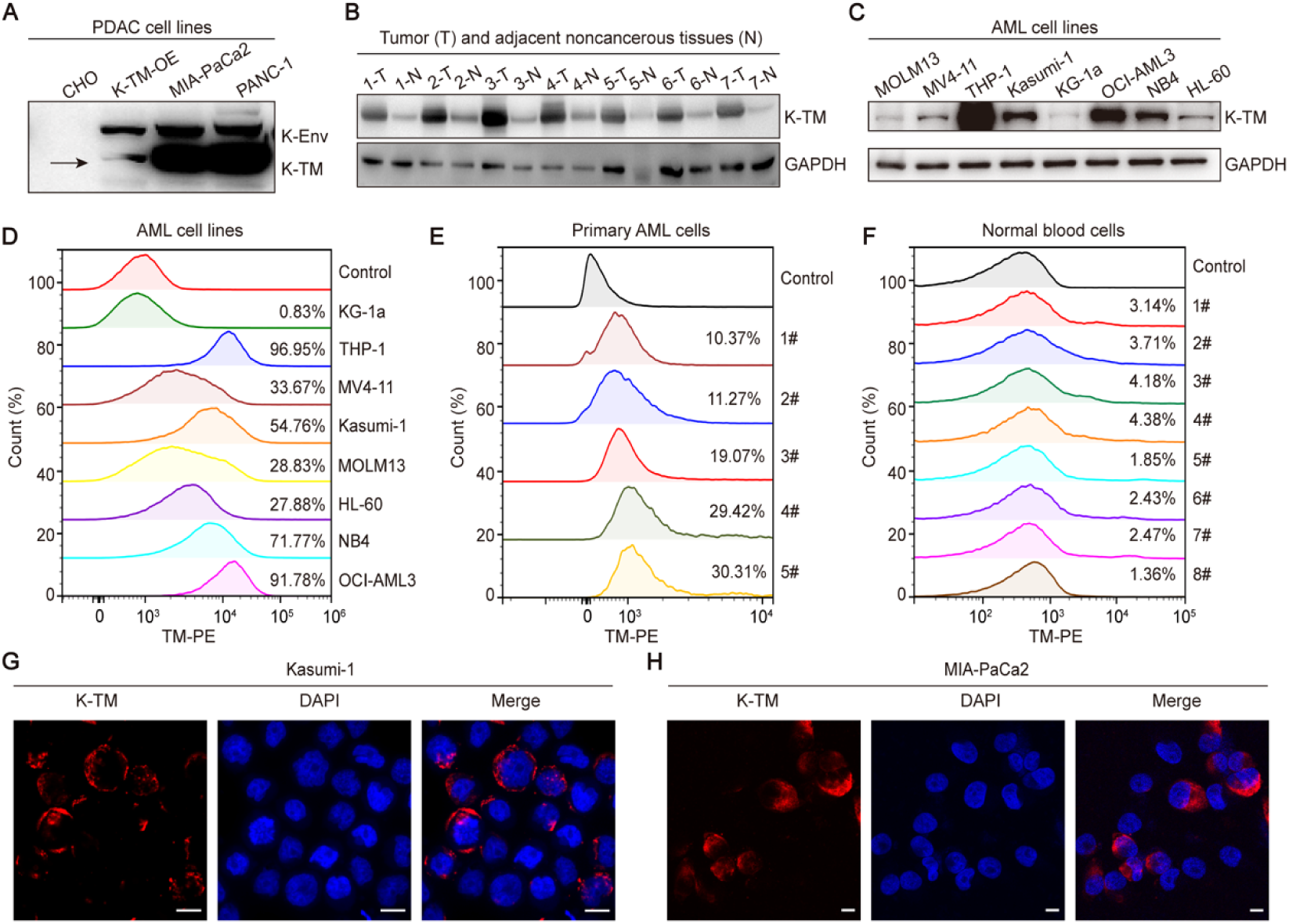
HERV-K102 Env-TM autoantibody-reactive antigens are present in cancer cells. (**A**) K-TM autoantibody-reactive antigens were detected in pancreatic cancer cell lines by Western blotting. (**B**) Expression levels of K-TM autoantibody-reactive antigens in 7 pairs of primary PDAC tumors and adjacent noncancerous tissues were evaluated by Western blotting. (**C**) Expression levels of K-TM autoantibody-reactive antigens were assessed in human AML cell lines by Western blotting. (**D, E, F**) AML cell lines, primary AML cells (n = 5) and normal blood cells (n = 8) were analyzed for K-TM autoantibody-reactive antigens by FCM using the 186 autoantibody and the secondary antibody PE F(ab’)2 Goat anti-human IgG Fcγ. PE Mouse IgG1, κ Isotype Ctrl (FC) was used as a control antibody. (**G, H**) Representative immunofluorescence images showing K-TM autoantibody-reactive antigens in Kasumi-1 and MIA-PaCa2 cells (scale bar, 10 µm).

To ascertain the presence of K-TM autoantibody-reactive antigens in blood cancer cells, we examined their expression levels in human AML cell lines using Western blotting. Consistent with the findings in PDAC, the results demonstrated that K-TM autoantibody-reactive antigens were detected in the AML cell lines Kasumi-1, THP-1 and OCI-AML3 (Fig. 6C).

To assess the presence of K-TM autoantibody-reactive antigens on the surface of cancer cells, FCM was used to detect the antigens in a panel of cancer cell lines. The autoantibodies specifically bound to the surface of cancer cells in human AML cell lines, with the positive rates for each type of AML cell presented in Fig. 6D. The quantity of K-TM autoantibody-reactive antigens on the surface of different AML cell types varied, with autoantibody binding rates ranging from 96.95% for THP-1 cells to only 0.83% for KG-1a cells. Subsequently, the expression of K-TM autoantibody-reactive antigens was examined in primary AML cells (Fig. 6E) and normal blood cells (Fig. 6F) by FCM. The results indicated that K-TM autoantibody-reactive antigens were expressed in all primary tumor tissues but were low or absent in normal blood cell samples.

To determine the distribution of K-TM autoantibody-reactive antigens in cancer cells, immunostaining was performed on Kasumi-1 cells (Fig. 6G) and MIA-PaCa2 cells (Fig. 6H) using mAb 186 and the secondary antibody PE F(ab’)2 Goat anti-human IgG Fcγ. The results revealed that K-TM autoantibody-reactive antigens were present both on the surface and in the cytoplasm of cancer cells. Representative immunostaining images for several other AML cell lines, including KG-1a, OCI-AML3, THP-1, and MV4-11, are shown in Supplementary Fig. 4A-D.

Collectively, these data suggest that AML and PDAC cells express a substantial quantity of K-TM autoantibody-reactive antigens.

### Autoantibodies induce ADCC activity against cancer cells

Given that HERV-K envelope-reactive antibodies from plasma of patients with LUAD exhibit ADCC against A549 cancer cells^21^, we next investigated whether autoantibodies from SLE patients induce ADCC activity against PDAC cancer cell lines and AML cell lines. ADCC assays were performed with 186 and 910 autoantibodies against Mia-PaCa2 cells expressing K-TM using healthy human PBMCs (n = 5) at an E:T ratio of 3:1, as depicted in Fig. 7A, B. ADCC killing effects for other autoantibodies are presented in Supplementary Fig.5A-I. Significant ADCC activity by autoantibodies (at 1 μg/mL and 5 μg/mL concentrations) was observed against the MIA-PaCa2 cells. Subsequently, we explored whether 186 and 910 autoantibodies induced ADCC effects on OCI-AML3 cells. PBMCs were isolated from four healthy individuals at an E:T ratio of 3:1, and the killing effect of autoantibodies on AML tumor cells was assessed by co-incubating them with OCI-AML3 cells using antibody concentrations of 1 μg/mL and 5 μg/mL (Fig. 7C, D). The results demonstrated that K-TM autoantibodies exhibited excellent ADCC effects not only on PDAC cell lines but also on AML cell lines. These findings suggest that autoantibodies from SLE patients could activate ADCC activity against both PDAC and AML cancer cells.

**Fig. 7.**
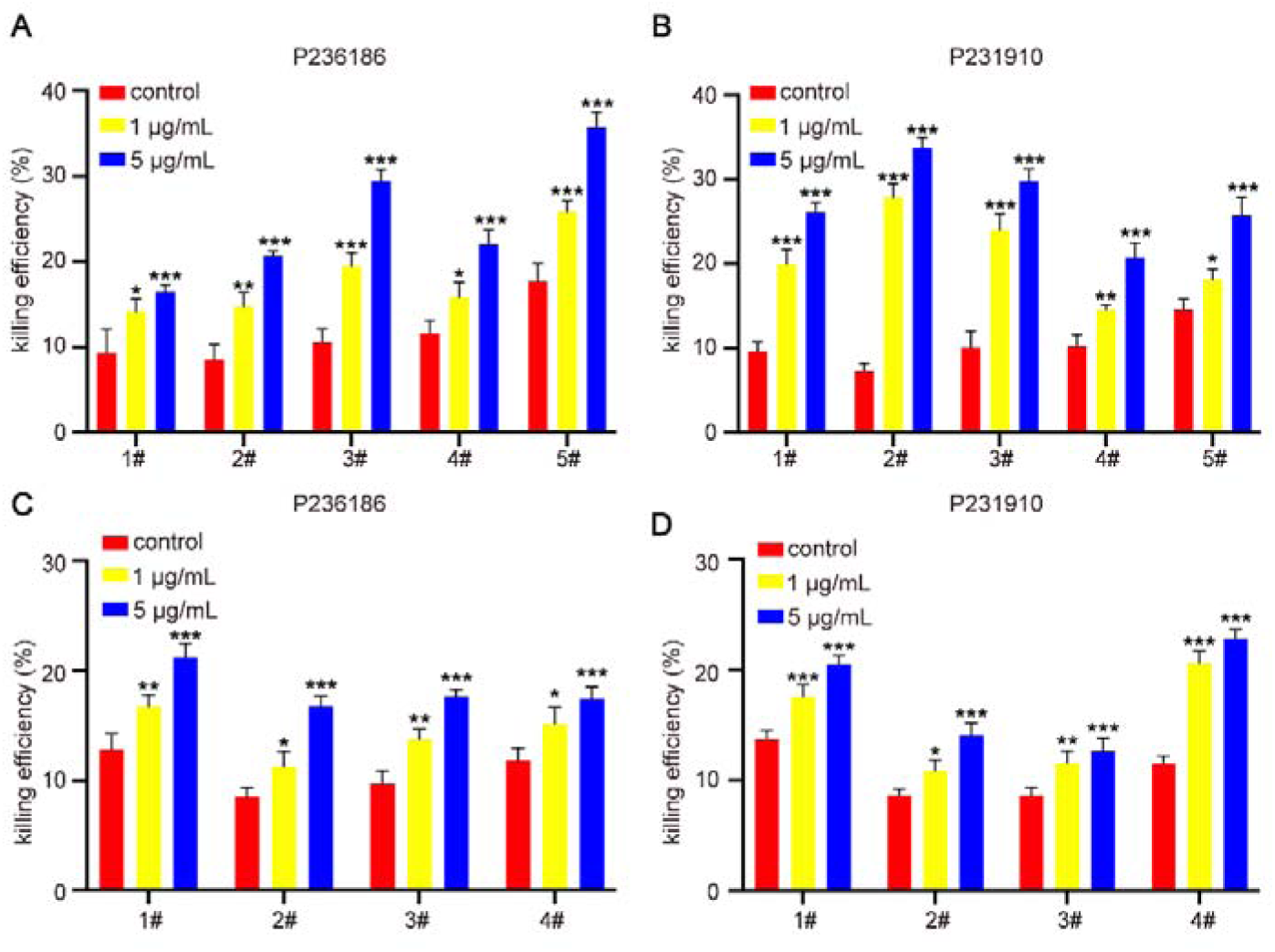
Autoantibodies induce ADCC activity against PDAC and AML cell lines. (**A, B**) An ADCC assay was performed using 186 and 910 autoantibodies targeting Mia-PaCa2 cells, with healthy human PBMCs serving as effector cells (n = 5). (**C, D**) An ADCC assay was conducted using 186 and 910 autoantibodies directed against OCI-AML3 cells, with healthy human PBMCs acting as effector cells (n = 4). Data are presented as mean values ± s.d. of 3 biological replicates. The differences between experimental groups and control group were compared, * p < 0.05, ** p < 0.01, *** p < 0.001, and ns p > 0.05.

## Discussion

We have developed a novel fully human anti-K102 Env-TM autoantibody phage display library with 3.67×10^8^ cfu using PBMCs from SLE patients and obtained 101 K-TM antigen-binding IgG1^+^ clonotypes and 48 K-SU antigen-binding IgG1^+^ clonotypes through high-throughput screening with HERV-K102-TM or SU antigens. We selected 135 of these clonotypes for sequencing analysis and obtained 68 unique molecules, from which 20 were chosen for full-length construction. Among the 20 full-length clones, 11 mAb clones only reacted with the K102-Env-TM glycoprotein, 8 mAb clones reacted with both K-SU and K-TM, and one mAb clone only reacted with K-SU. The EC50 values of these autoantibodies for binding to the TM subunit ranged from 0.002436 μg/mL to 1.798 μg/mL. These findings suggest that the in vitro binding affinity of monoclonal autoantibodies against the HERV-K102 Env-TM subunit antigen is well reflected in vivo in SLE patients.

Unexpectedly, we found high levels of HERV-K102 Env-TM^+^ B, T and NK cells in patients with active SLE. The positive rates of HERV-K102 Env-TM^+^ B, T and NK cells in one SLE patient were 52.16%, 31.45%, and 48.89%, respectively. In contrast, the positive rates of HERV-K102 Env-TM^+^ B, T and NK cells in one healthy control were 9.88%, 2.12%, and 10.76%, respectively. Furthermore, HERV-K102 Env-TM^+^ T cells were positive for IFN-α, and HERV-K102 Env-TM^+^ NK cells were positive for GZMB, indicating that these HERV-K102-Env-TM^+^ cells might be autoreactive immune cells.

Notably, our study demonstrated that HERV-K102 Env-TM monoclonal autoantibodies selectively eliminate HERV-K102 Env-TM^+^ immune cells while exhibiting no impact on normal immune cells. These findings suggest that these autoantibodies may contribute to the normalization of immune cell subsets in SLE patients.

This study reveals that AML and PDAC cancer cells express a significant quantity of K-TM autoantibody-reactive antigens. K-TM monoclonal autoantibodies not only recognize K-TM glycoprotein on the cell surface, but also exhibit potent activity for eliminating cancer cells. Crucially, the autoantibody-reactive K-TM antigens are localized on the surface of cancer cells, rendering them potential targets for therapeutic intervention.

In summary, we established the world’s first fully human autoantibody phage display library with high capacity and identified a series of mAbs exhibiting potent activity against the conserved HERV-K102 Env-TM subunit. These monoclonal autoantibodies not only recognize the HERV-K102 Env-TM glycoprotein on the cell surface but also eliminate autoreactive B, T and NK cells in SLE as well as cancer cells. Our findings provide a conceptually novel immunotherapy for autoimmune diseases and cancers by targeting the HERV-K102 Env-TM subunit and open the era of using human autoantibodies to treat autoimmune diseases and cancers. In addition, several of these monoclonal autoantibodies are promising candidates for clinical development as potential diagnostic and therapeutic agents, paving the way for treating SLE and cancers. Autoantibodies from patients with autoimmune diseases can be used as natural biologic drugs with an extraordinary array of potential impacts on human cancers and autoimmune diseases.

## Acknowledgments

This work was supported by the National Natural Science Foundation of China (82150116, 82070146, and 82200163), and the Natural Science Foundation of Zhejiang Province (LZ21H160005 and LQ22H080007).

## Ethics approval statement

This study protocol was approved by the Ethics and Scientific Committee of The Second Affiliated Hospital of Zhejiang University School of Medicine (No.2024–0325).

## Author contributions

R.Z.X conceived the study, initiated, designed, and supervised the experiments. R.Z.X, Q.Y.G and M.Y.L wrote the manuscript. Q.Y.G, M.Y.L, S.W.Z, Z.X.W, and X.Z.Z performed experiments. Y.L, W.B.Q, and R.Z.X supervised the study and experiments.

## Data availability statement

The data that support the findings of this study are available from the corresponding author upon reasonable request.

---

**Supplementary Fig. 1.**
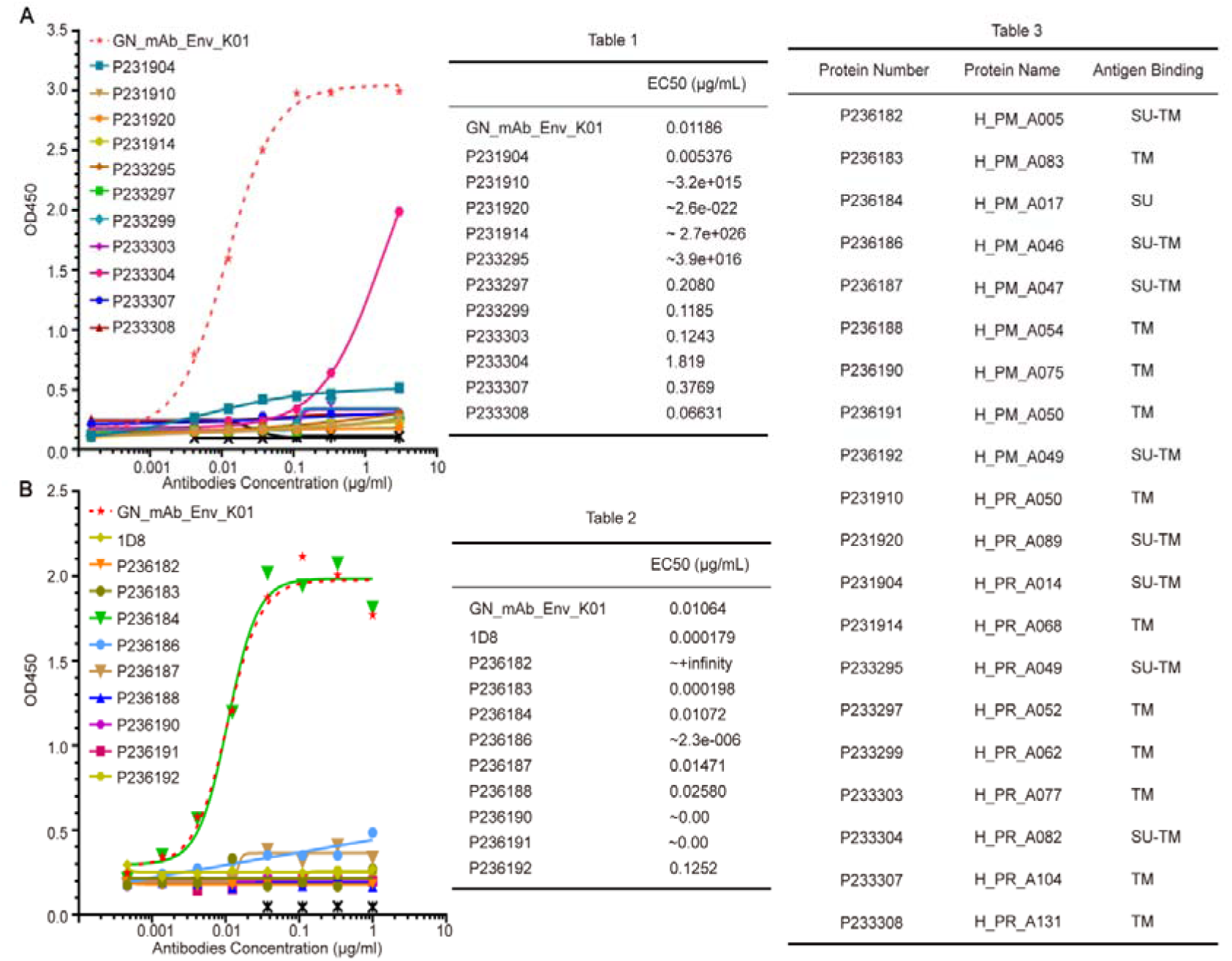
Binding affinity of 20 candidate antibodies to K102-SU-FC antigen by ELISA. (**A**) 904, 910, 920, 914, 295, 297, 299, 303, 304, 307 and 308 antibodies. (**B**) 182, 183, 184, 186, 187, 188, 190, 191 and 192 antibodies.

**Supplementary Fig. 2.**
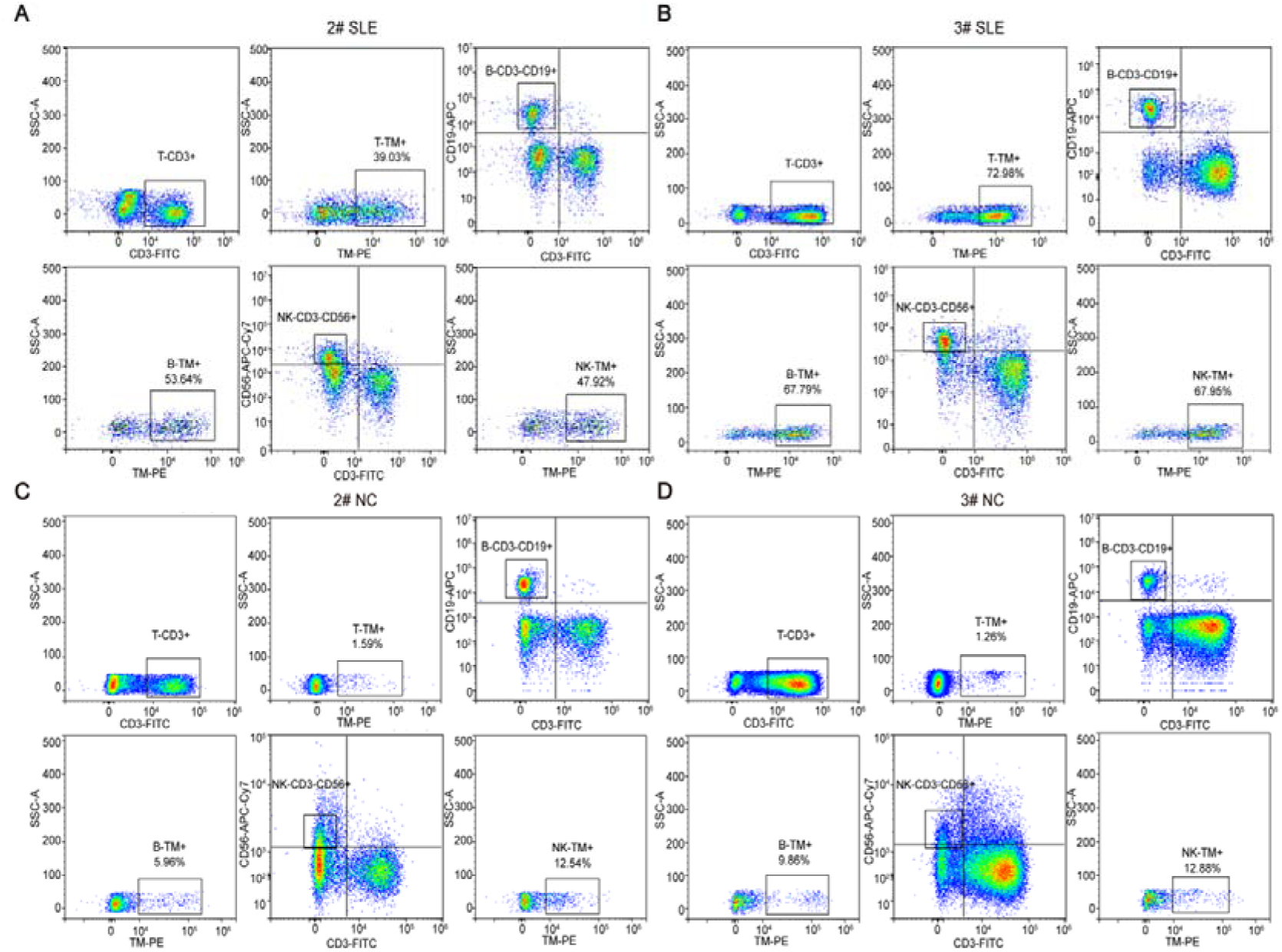
The proportions of HERV-K102 Env-TM^+^ T, B and NK cells in the blood of SLE patients and healthy controls. (A) FC plots and gating strategies employed to identify K-TM^+^ immune cells in 2# and 3# SLE patients. (**B**) FC plots and gating strategies utilized to identify K-TM^+^ immune cells in 2# and 3# healthy controls.

**Supplementary Fig. 3.**
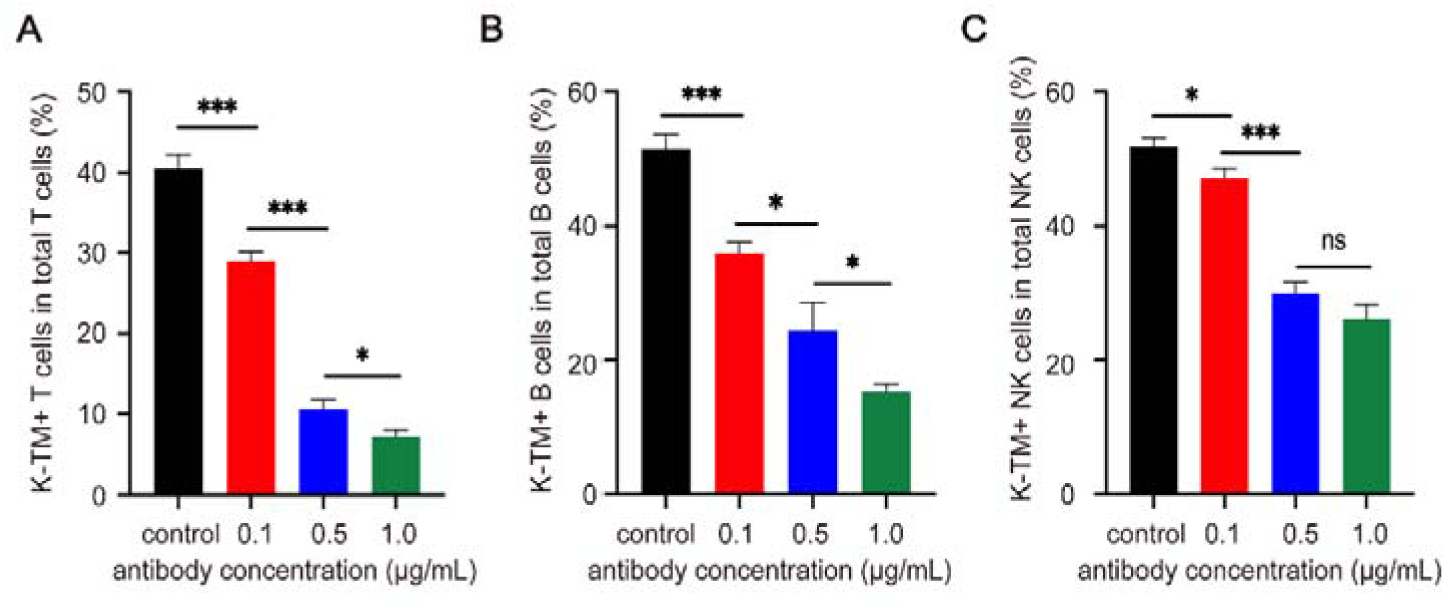
The ADCC activity of HERV-K102 Env-TM monoclonal autoantibody to HERV-K102 Env-TM^+^ T, B and NK cells in 2# active SLE patient. (**A-C**) Various concentrations of the autoantibody (0.1 μg/mL, 0.5 μg/mL and 1 μg/mL) eliminated K-TM^+^ T, B, and NK cells from SLE patients. Data are presented as mean values ± s.d. of 3 biological replicates. * p < 0.05, ** p < 0.01, *** p < 0.001, and ns p > 0.05.

**Supplementary Fig. 4.**
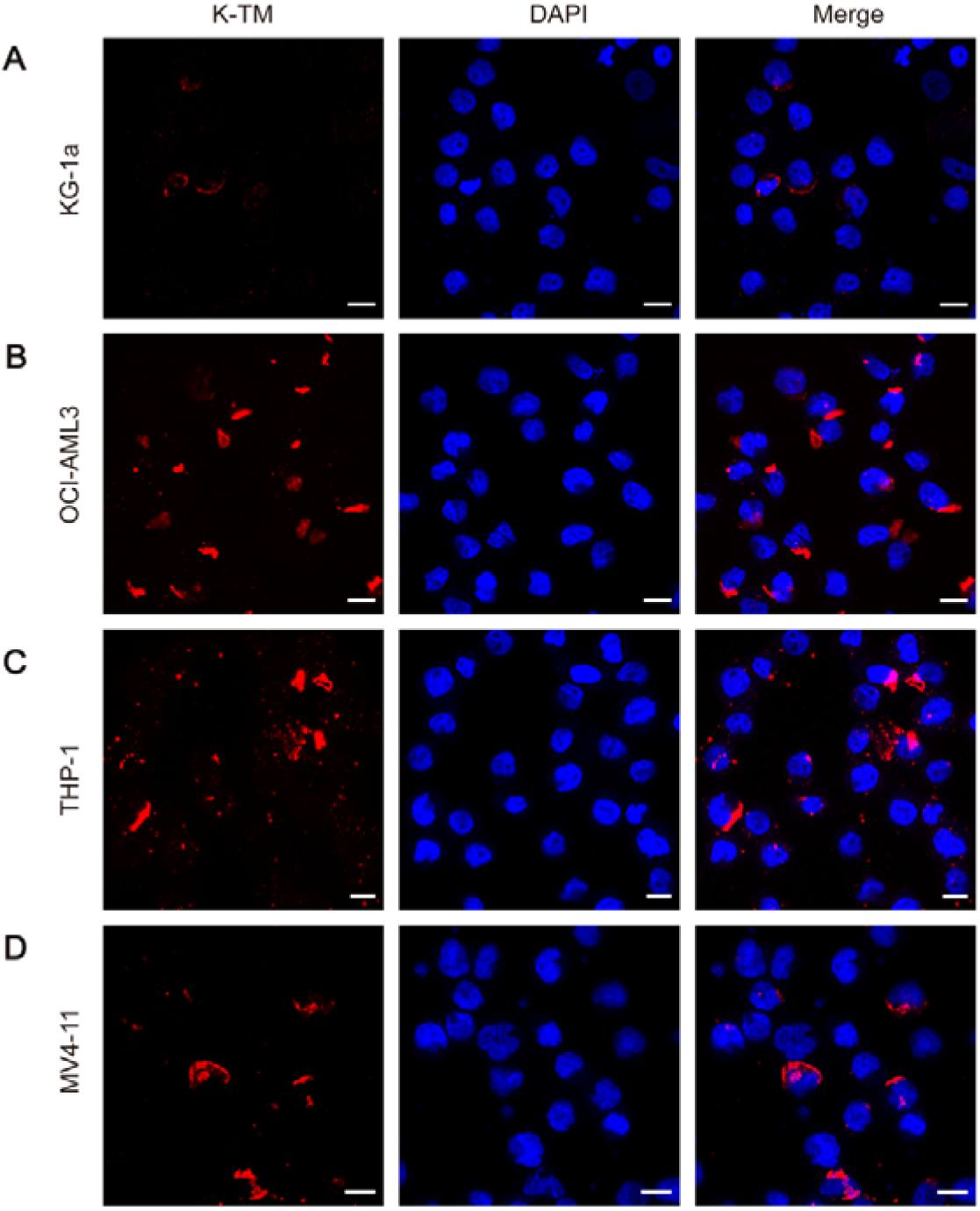
HERV-K102 Env-TM autoantibody-reactive antigens present in AML cell lines. (**A-D**) Representative immunofluorescence images illustrating the presence of K-TM autoantibody-reactive antigens in KG-1a, OCI-AML3, THP-1 and MV4-11 cells (scale bar, 10 µm).

**Supplementary Fig. 5.**
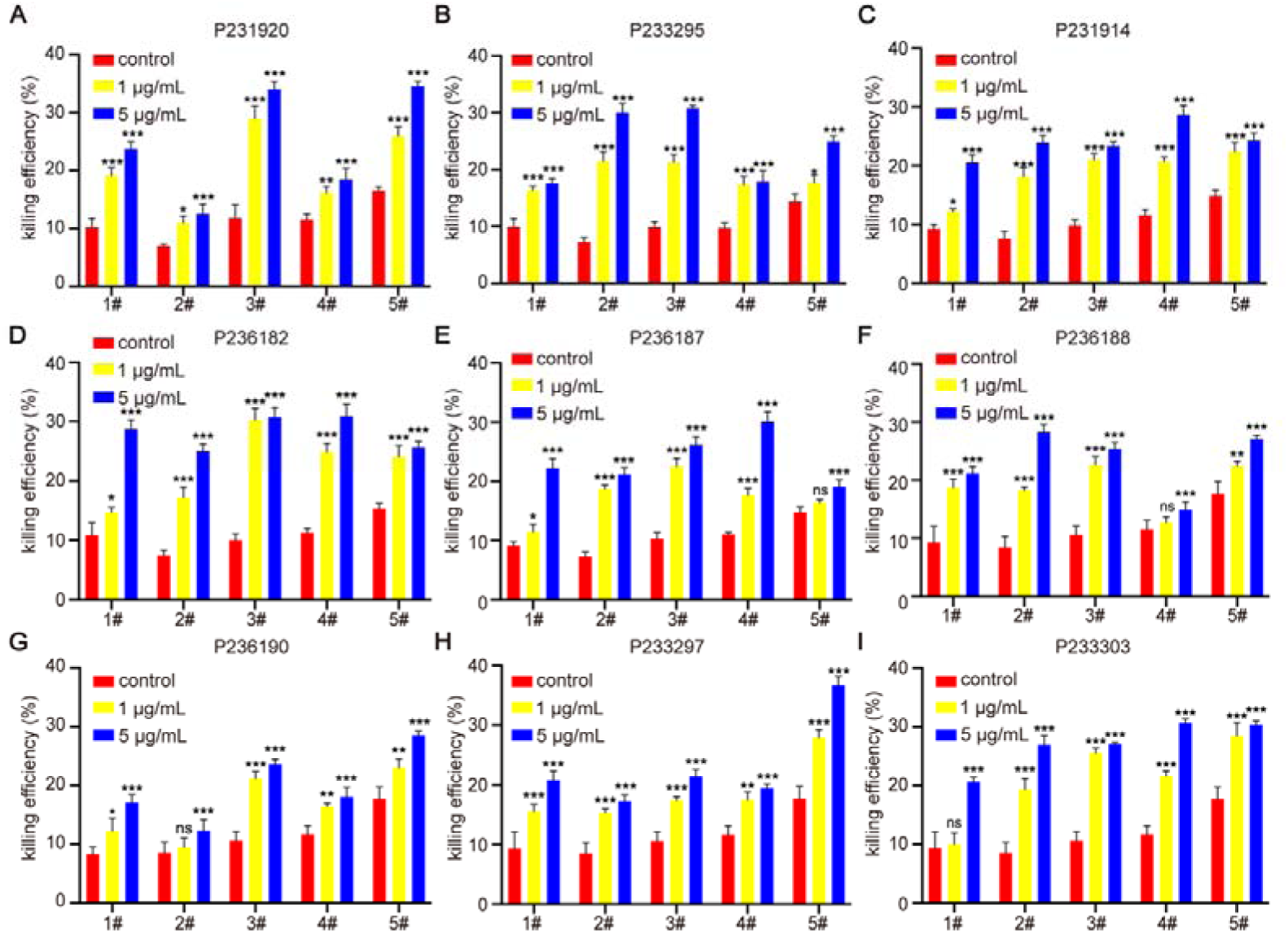
Autoantibodies induce ADCC activity against MIA-PaCa2 cells. The 920 (**A**), 295 (**B**), 914 (**C**), 182 (**D**), 187 (**E**), 188 (**F**), 190 (**G**), 297 (**H**), and 303 (**I**) antibodies. Data are presented as mean values ± s.d. of 3 biological replicates. The differences between experimental groups and control group were compared, * p < 0.05, ** p < 0.01, *** p < 0.001, and ns p > 0.05.

**Supplementary** Table 1 EC50 values of 904, 910, 920, 914, 295, 297, 299, 303, 304, 307 and 308 antibodies.

**Supplementary** Table 2 EC50 values of 182, 183, 184, 186, 187, 188, 190, 191 and 192 antibodies.

**Supplementary** Table 3 Binding assay of 20 mAbs to K-TM and K-SU antigens.

## Reference

1. Chen D. S., Mellman I. Elements of cancer immunity and the cancer-immune set point. Nature. (2017) 541:321–330. 10.1038/nature21349

2. Li J., Wu C., Hu H., Qin G., Wu X., Bai F., et al. Remodeling of the immune and stromal cell compartment by PD-1 blockade in mismatch repair-deficient colorectal cancer. Cancer Cell. (2023) 41:1152–1169.e1157. 10.1016/j.ccell.2023.04.011

3. Hawley J. E., Obradovic A. Z., Dallos M. C., Lim E. A., Runcie K., Ager C. R., et al. Anti-PD-1 immunotherapy with androgen deprivation therapy induces robust immune infiltration in metastatic castration-sensitive prostate cancer. Cancer Cell. (2023) 41:1972–1988.e1975. 10.1016/j.ccell.2023.10.006

4. Tousley A. M., Rotiroti M. C., Labanieh L., Rysavy L. W., Kim W. J., Lareau C., et al. Co-opting signalling molecules enables logic-gated control of CAR T cells. Nature. (2023) 615:507–516. 10.1038/s41586-023-05778-2

5. Raje N., Berdeja J., Lin Y., Siegel D., Jagannath S., Madduri D., et al. Anti-BCMA CAR T-Cell Therapy bb2121 in Relapsed or Refractory Multiple Myeloma. N Engl J Med. (2019) 380:1726–1737. 10.1056/NEJMoa1817226

6. Haradhvala N. J., Leick M. B., Maurer K., Gohil S. H., Larson R. C., Yao N., et al. Distinct cellular dynamics associated with response to CAR-T therapy for refractory B cell lymphoma. Nat Med. (2022) 28:1848–1859. 10.1038/s41591-022-01959-0

7. Mackensen A., Müller F., Mougiakakos D., Böltz S., Wilhelm A., Aigner M., et al. Anti-CD19 CAR T cell therapy for refractory systemic lupus erythematosus. Nat Med. (2022) 28:2124–2132. 10.1038/s41591-022-02017-5

8. Müller F., Taubmann J., Bucci L., Wilhelm A., Bergmann C., Völkl S., et al. CD19 CAR T-Cell Therapy in Autoimmune Disease - A Case Series with Follow-up. N Engl J Med. (2024) 390:687–700. 10.1056/NEJMoa2308917

9. Sosnoski H. M., Posey A. D., Jr. Therapeutic intersections: Expanding benefits of CD19 CAR T cells from cancer to autoimmunity. Cell Stem Cell. (2024) 31:437–438. 10.1016/j.stem.2024.03.006

10. O’Donnell J. S., Teng M. W. L., Smyth M. J. Cancer immunoediting and resistance to T cell-based immunotherapy. Nat Rev Clin Oncol. (2019) 16:151–167. 10.1038/s41571-018-0142-8

11. Hegde P. S., Chen D. S. Top 10 Challenges in Cancer Immunotherapy. Immunity. (2020) 52:17–35. 10.1016/j.immuni.2019.12.011

12. Chow A., Perica K., Klebanoff C. A., Wolchok J. D. Clinical implications of T cell exhaustion for cancer immunotherapy. Nat Rev Clin Oncol. (2022) 19:775–790. 10.1038/s41571-022-00689-z

13. Yousuf S., Qiu M., Voith von Voithenberg L., Hulkkonen J., Macinkovic I., Schulz A. R., et al. Spatially Resolved Multi-Omics Single-Cell Analyses Inform Mechanisms of Immune Dysfunction in Pancreatic Cancer. Gastroenterology. (2023) 165:891–908.e814. 10.1053/j.gastro.2023.05.036

14. Rutella S., Vadakekolathu J., Mazziotta F., Reeder S., Yau T. O., Mukhopadhyay R., et al. Immune dysfunction signatures predict outcomes and define checkpoint blockade-unresponsive microenvironments in acute myeloid leukemia. J Clin Invest. (2022) 132:10.1172/jci159579

15. Qiu J., Xu B., Ye D., Ren D., Wang S., Benci J. L., et al. Cancer cells resistant to immune checkpoint blockade acquire interferon-associated epigenetic memory to sustain T cell dysfunction. Nat Cancer. (2023) 4:43–61. 10.1038/s43018-022-00490-y

16. Khadjinova A. I., Wang X., Laine A., Ukadike K., Eckert M., Stevens A., et al. Autoantibodies against the envelope proteins of endogenous retroviruses K102 and K108 in patients with systemic lupus erythematosus correlate with active disease. Clin Exp Rheumatol. (2022) 40:1306–1312. 10.55563/clinexprheumatol/2kg1d8

17. Tokuyama M., Gunn B. M., Venkataraman A., Kong Y., Kang I., Rakib T., et al. Antibodies against human endogenous retrovirus K102 envelope activate neutrophils in systemic lupus erythematosus. J Exp Med. (2021) 218:10.1084/jem.20191766

18. Law C., Wacleche V. S., Cao Y., Pillai A., Sowerby J., Hancock B., et al. Interferon subverts an AHR-JUN axis to promote CXCL13(+) T cells in lupus. Nature. (2024) 631:857–866. 10.1038/s41586-024-07627-2

19. Mori S., Kohyama M., Yasumizu Y., Tada A., Tanzawa K., Shishido T., et al. Neoself-antigens are the primary target for autoreactive T cells in human lupus. Cell. (2024) 10.1016/j.cell.2024.08.025

20. Chen T., Meng Z., Gan Y., Wang X., Xu F., Gu Y., et al. The viral oncogene Np9 acts as a critical molecular switch for co-activating β-catenin, ERK, Akt and Notch1 and promoting the growth of human leukemia stem/progenitor cells. Leukemia. (2013) 27:1469–1478. 10.1038/leu.2013.8

21. Ng K. W., Boumelha J., Enfield K. S. S., Almagro J., Cha H., Pich O., et al. Antibodies against endogenous retroviruses promote lung cancer immunotherapy. Nature. (2023) 616:563–573. 10.1038/s41586-023-05771-9

22. Shah A. H., Rivas S. R., Doucet-O’Hare T. T., Govindarajan V., DeMarino C., Wang T., et al. Human endogenous retrovirus K contributes to a stem cell niche in glioblastoma. J Clin Invest. (2023) 133:10.1172/jci167929

23. Camargo-Forero N., Orozco-Arias S., Perez Agudelo J. M., Guyot R. HERV-K (HML-2) insertion polymorphisms in the 8q24.13 region and their potential etiological associations with acute myeloid leukemia. Arch Virol. (2023) 168:125. 10.1007/s00705-023-05747-0

24. Deniz Ö, Ahmed M., Todd C. D., Rio-Machin A., Dawson M. A., Branco M. R. Endogenous retroviruses are a source of enhancers with oncogenic potential in acute myeloid leukaemia. Nat Commun. (2020) 11:3506. 10.1038/s41467-020-17206-4

25. Li M., Radvanyi L., Yin B., Rycaj K., Li J., Chivukula R., et al. Downregulation of Human Endogenous Retrovirus Type K (HERV-K) Viral env RNA in Pancreatic Cancer Cells Decreases Cell Proliferation and Tumor Growth. Clin Cancer Res. (2017) 23:5892–5911. 10.1158/1078-0432.Ccr-17-0001

26. Xue B., Sechi L. A., Kelvin D. J. Human Endogenous Retrovirus K (HML-2) in Health and Disease. Front Microbiol. (2020) 11:1690. 10.3389/fmicb.2020.01690

27. Bradford H. F., Haljasmägi L., Menon M., McDonnell T. C. R., Särekannu K., Vanker M., et al. Inactive disease in patients with lupus is linked to autoantibodies to type I interferons that normalize blood IFNα and B cell subsets. Cell Rep Med. (2023) 4:100894. 10.1016/j.xcrm.2022.100894

28. Demers-Mathieu V. Optimal Selection of IFN-α-Inducible Genes to Determine Type I Interferon Signature Improves the Diagnosis of Systemic Lupus Erythematosus. Biomedicines. (2023) 11:10.3390/biomedicines11030864

29. Chaudhary V., Ah Kioon M. D., Hwang S. M., Mishra B., Lakin K., Kirou K. A., et al. Chronic activation of pDCs in autoimmunity is linked to dysregulated ER stress and metabolic responses. J Exp Med. (2022) 219:10.1084/jem.20221085

30. Rahaman O., Bhattacharya R., Liu C. S. C., Raychaudhuri D., Ghosh A. R., Bandopadhyay P., et al. Cutting Edge: Dysregulated Endocannabinoid-Rheostat for Plasmacytoid Dendritic Cell Activation in a Systemic Lupus Endophenotype. J Immunol. (2019) 202:1674–1679. 10.4049/jimmunol.1801521

31. Sakata K., Nakayamada S., Miyazaki Y., Kubo S., Ishii A., Nakano K., et al. Up-Regulation of TLR7-Mediated IFN-α Production by Plasmacytoid Dendritic Cells in Patients With Systemic Lupus Erythematosus. Front Immunol. (2018) 9:1957. 10.3389/fimmu.2018.01957

32. Menon M., Blair P. A., Isenberg D. A., Mauri C. A Regulatory Feedback between Plasmacytoid Dendritic Cells and Regulatory B Cells Is Aberrant in Systemic Lupus Erythematosus. Immunity. (2016) 44:683–697. 10.1016/j.immuni.2016.02.012

