## supplemental file for "Natural monoclonal autoantibodies against HERV-K102 Envelope-TM from SLE patients selectively eliminate autoreactive immune cells and cancer cells"

**Supplementary informationion**
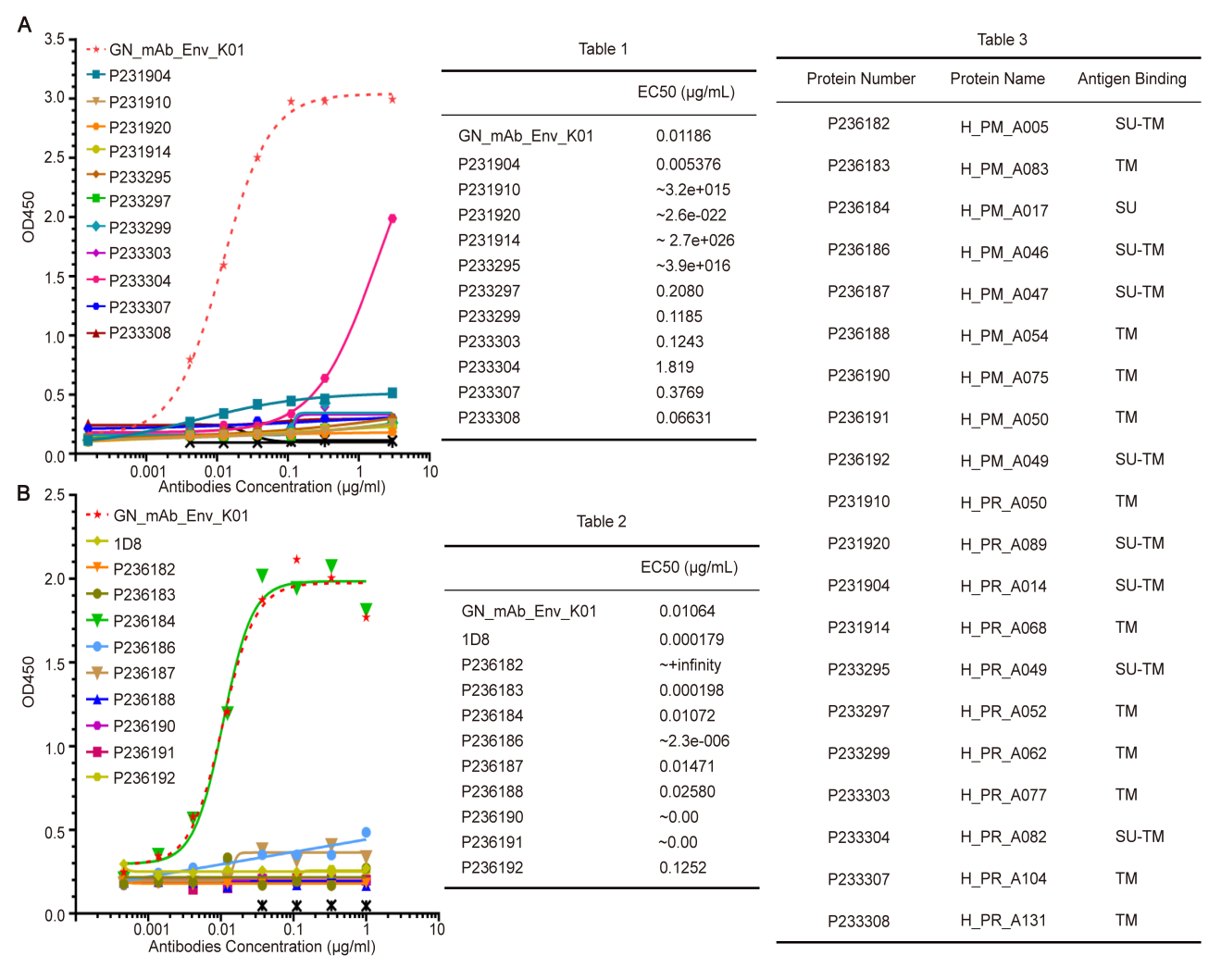

**Supplementary Fig. 1** Binding affinity of 20 candidate antibodies to K102-SU-FC antigen by ELISA. (**A**) 904, 910, 920, 914, 295, 297, 299, 303, 304, 307 and 308 antibodies. (**B**) 182, 183, 184, 186, 187, 188, 190, 191 and 192 antibodies.

**Supplementary Table 1** EC50 values of 904, 910, 920, 914, 295, 297, 299, 303, 304, 307 and 308 antibodies.

**Supplementary Table 2** EC50 values of 182, 183, 184, 186, 187, 188, 190, 191 and 192 antibodies.

**Supplementary Table 3** Binding assay of 20 mAbs to K-TM and K-SU antigens.

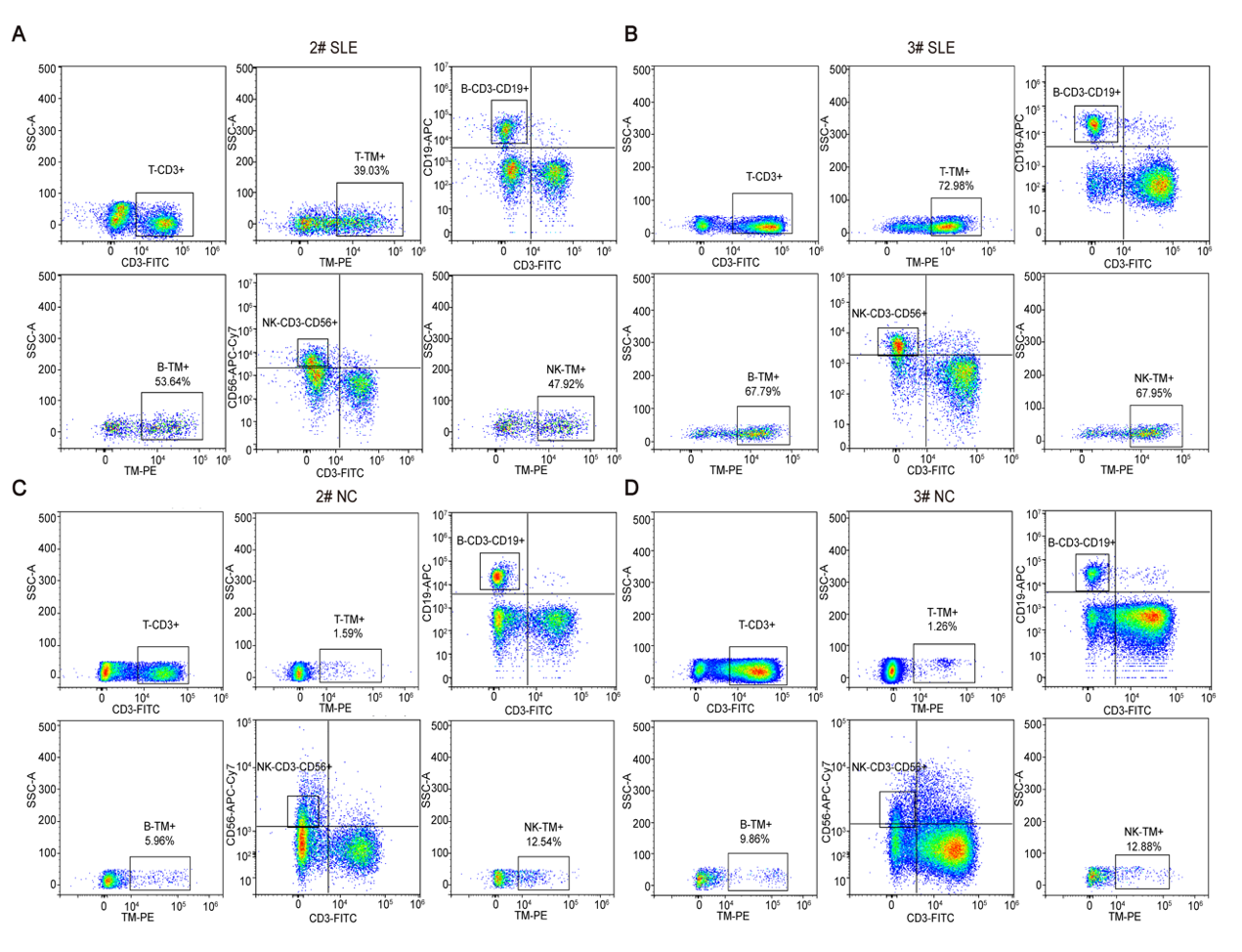

**Supplementary Fig. 2** The proportions of HERV-K102 Env-TM^+^ T, B and NK cells in the blood of SLE patients and healthy controls.

**(**A) FC plots and gating strategies employed to identify K-TM^+^ immune cells in 2# and 3# SLE patients. (**B**) FC plots and gating strategies utilized to identify K-TM^+^ immune cells in 2# and 3# healthy controls.

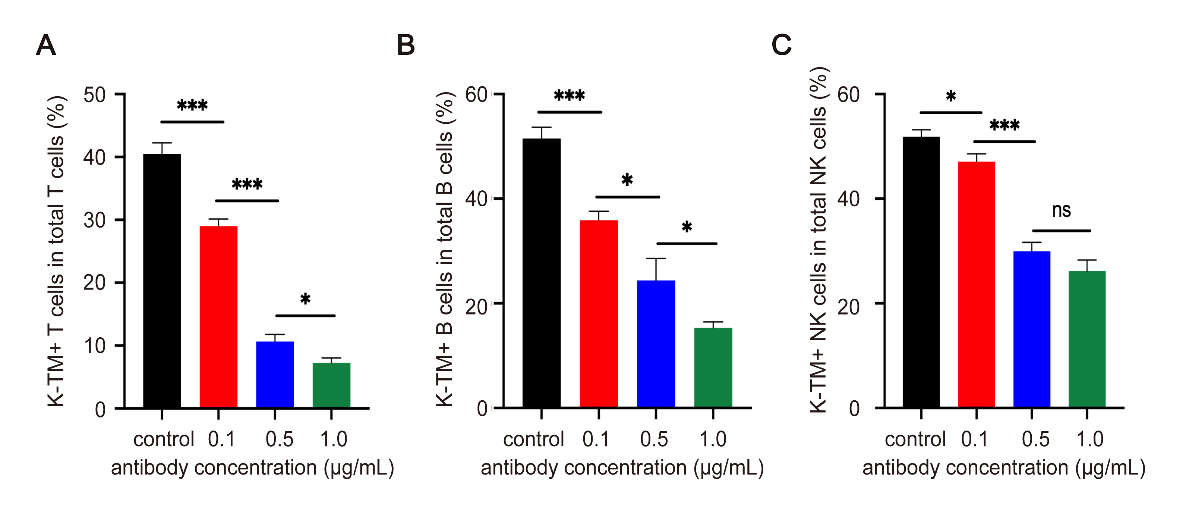

**Supplementary Fig. 3** The ADCC activity of HERV-K102 Env-TM monoclonal autoantibody to HERV-K102 Env-TM^+^ T, B and NK cells in 2# active SLE patient. (**A-C**) Various concentrations of the autoantibody (0.1 μg/mL, 0.5 μg/mL and 1 μg/mL) eliminated K-TM^+^ T, B, and NK cells from SLE patients. Data are presented as mean values ± s.d. of 3 biological replicates. * p < 0.05, ** p < 0.01, *** p < 0.001, and ns p > 0.05.

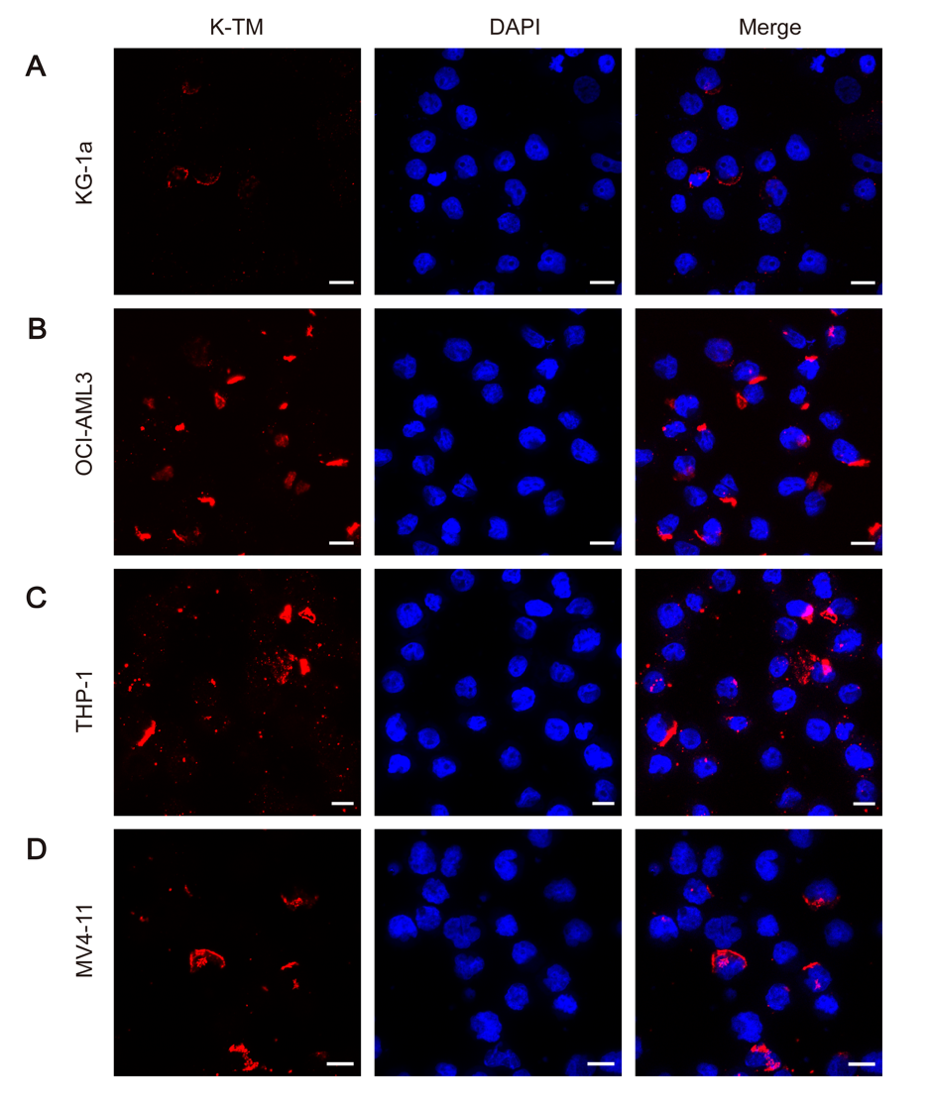

**Supplementary Fig. 4** HERV-K102 Env-TM autoantibody-reactive antigens present in AML cell lines. (**A-D**) Representative immunofluorescence images illustrating the presence of K-TM autoantibody-reactive antigens in KG-1a, OCI-AML3, THP-1 and MV4-11 cells (scale bar, 10 µm).

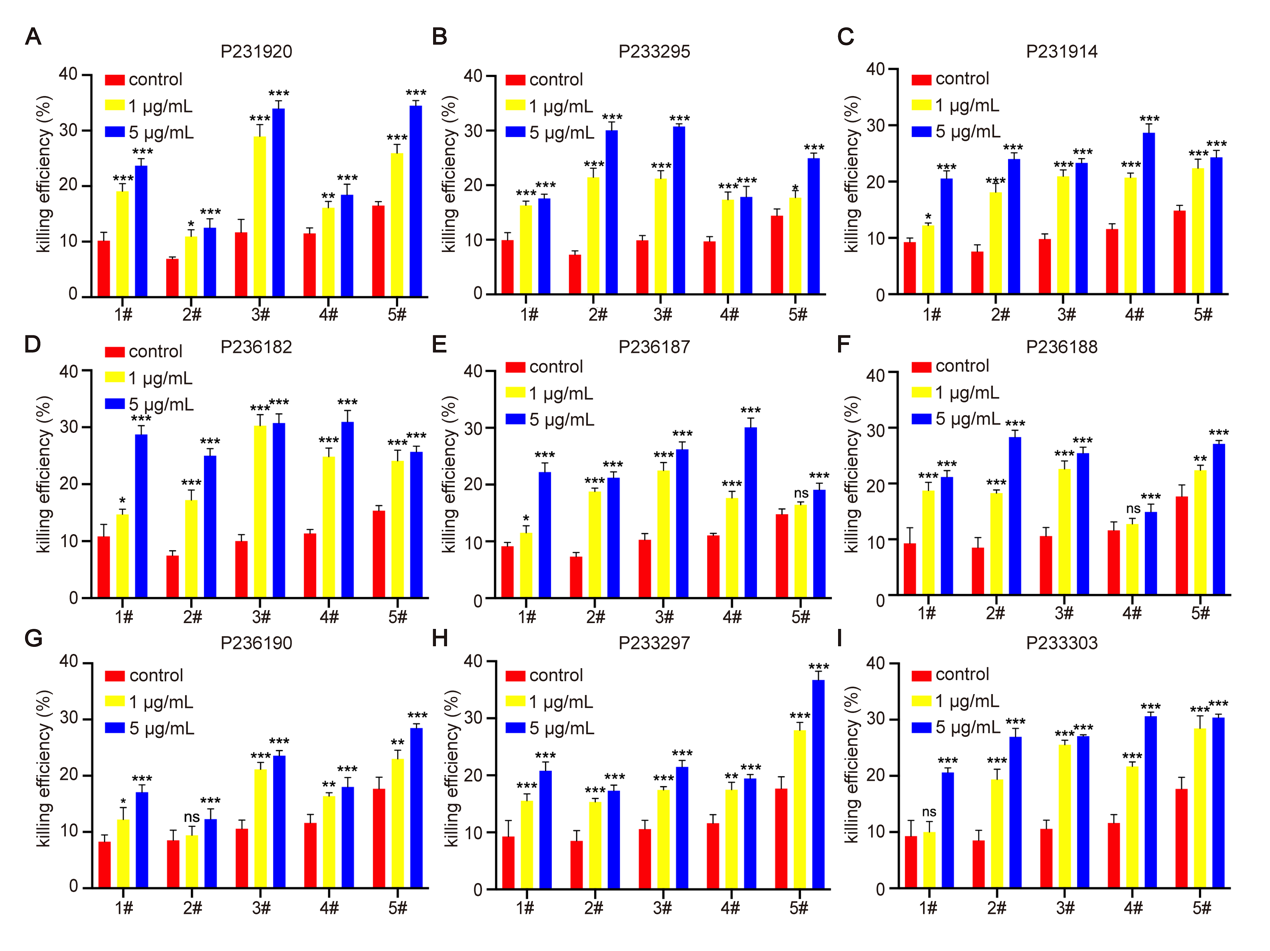

**Supplementary Fig. 5** Autoantibodies induce ADCC activity against MIA-PaCa2 cells. The 920 (**A**), 295 (**B**), 914 (**C**), 182 (**D**), 187 (**E**), 188 (**F**), 190 (**G**), 297 (**H**), and 303 (**I**) antibodies. Data are presented as mean values ± s.d. of 3 biological replicates. The differences between experimental groups and control group were compared, * p < 0.05, ** p < 0.01, *** p < 0.001, and ns p > 0.05.
